# Synthesis of a eukaryotic chromosome reveals a role for N6-methyladenine in nucleosome organization

**DOI:** 10.1101/184929

**Authors:** Leslie Y. Beh, Galia T. Debelouchina, Kelsi A. Lindblad, Katarzyna Kulej, Elizabeth R. Hutton, John R. Bracht, Robert P. Sebra, Benjamin A. Garcia, Tom W. Muir, Laura F. Landweber

## Abstract

Biochemical studies of chromatin have typically used either artificial DNA templates with unnaturally high affinity for histones, or small genomic DNA fragments deprived of their cognate physical environment. It has thus been difficult to dissect chromatin structure and function within fully native DNA substrates. Here, we circumvent these limitations by exploiting the minimalist genome of the eukaryote *Oxytricha trifallax*, whose notably small ~3kb chromosomes mainly encode single genes. Guided by high-resolution epigenomic maps of nucleosome organization, transcription, and DNA N^6^-methyladenine (m^6^dA) locations, we reconstruct full-length *Oxytricha* chromosomes *in vitro* and use these synthetic facsimiles to dissect the influence of m^6^dA and histone post-translational modifications on nucleosome organization. We show that m^6^dA directly disfavors nucleosomes in a quantitative manner, leading to local decreases in nucleosome occupancy that are synergistic with histone acetylation. The effect of m^6^dA can be partially reversed by the action of an ATP-dependent chromatin remodeler. Furthermore, erasing m^6^dA marks from *Oxytricha* chromosomes leads to proportional increases in nucleosome occupancy across the genome. This work showcases *Oxytricha* chromosomes as powerful yet practical models for studying eukaryotic chromatin and transcription in the context of biologically relevant DNA substrates.

**Highlights:**

- *De novo* synthesis of complete, epigenetically defined *Oxytricha* chromosomes
- Epigenomic profiles of chromatin organization in *Oxytricha’s* miniature chromosomes
- m^6^dA directly disfavors nucleosome occupancy in natural and synthetic chromosomes
- Histone acetylation and chromatin remodelers temper the impact of m^6^dA on chromatin

## Introduction

Nucleosomes are the fundamental repeating unit of eukaryotic chromatin, consisting of ~146bp DNA wrapped around an octameric core of histones. Nucleosomes limit the physical accessibility of DNA to trans-acting factors, and thus directly impact DNA-based transactions, such as transcription, DNA repair, and replication. The *in vitro* reconstitution of chromatin is a method central to understanding these processes at the molecular level. Currently, the most widely used DNA template for such experiments is the 147bp “601” nucleosome positioning sequence. It exhibits a ~374 fold higher affinity for histone octamers than native genomic sequences (Lowary and Widom, 1998; Thåström et al., 2004), allowing the preparation of consistent, defined nucleosome arrays *in vitro*. Yet the biological relevance of 601 DNA remains unclear, due to its unnaturally high affinity for histones. On the other hand, reconstituting chromatin from native genomic DNA is challenging because the long length of chromosomes increases the propensity for non-specific aggregation during chromatin assembly (Kaplan et al., 2009). As a compromise, small genomic fragments are usually prepared via PCR amplification or mechanical shearing. However, such substrates are separated from their cognate physical environment, as defined by the totality of the chromosome.

Eukaryotic genomes exhibit enormous natural variation in form and function. This is exemplified by unicellular ciliates, which possess two structurally and functionally distinct nuclei within each cell (Prescott, 1994; Yerlici and Landweber, 2014). In *Oxytricha trifallax*, the **germline micronucleus** is transcriptionally silent and contains ~100 megabase-sized chromosomes, similar to widely studied model organisms (Chen et al., 2014). In contrast, the **somatic macronucleus** is transcriptionally active, being the sole locus of Pol II-dependent RNA production in non-developing cells (Khurana et al., 2014). The *Oxytricha* macronuclear genome is extraordinarily fragmented, consisting of ~16,000 unique chromosomes with a mean length of ~3.2kb, most encoding a single gene. Chromosome ends are capped with a compact 36nt telomere consisting of 5’-(T_4_G_4_)-3’ repeats, bound cooperatively by a heterodimeric protein complex (Gottschling and Zakian, 1986; Horvath et al., 1998). Macronuclear chromatin yields a characteristic ~200bp ladder upon digestion with micrococcal nuclease, indicative of regularly spaced nucleosomes (Gottschling and Cech, 1984; Lawn et al., 1978; Wada and Spear, 1980). Yet it remains unknown how and where nucleosomes are organized within these miniature chromosomes, and how transcriptional control is orchestrated in their context. Chromatin organization in *Oxytricha* offers a model system to shed light on these fundamental questions, and also opens exciting possibilities for constructing complete chromosomes with defined molecular composition. Such ‘designer’ chromosomes can allow investigation of histone and DNA modifications in the context of fully native DNA.

Ciliates have long served as powerful models for the study of chromatin modifications (Brownell et al., 1996; Liu et al., 2007; Strahl et al., 1999; Taverna et al., 2002; Wei et al., 1998). They also hold promise for the study of DNA methylation - in particular, N^6^-methyladenine (m^6^dA). This modification has recently been implicated in diverse biological processes in eukaryotes, including retrotransposon regulation, transgenerational epigenetic inheritance, and gene activation. m^6^dA is abundant in the macronuclear genomes of ciliates (0.18 - 2.5% m^6^dA / dA) (Ammermann et al., 1981; Cummings et al., 1974; Gorovsky et al., 1973; Rae and Spear, 1978), similar to the green algae *Chlamydomonas* (0.3 - 0.5%) (Fu et al., 2015; Hattman et al., 1978). High levels of m^6^dA (up to 2.8%) were also recently reported in basal fungi (Mondo et al., 2017). m^6^dA is present at very low levels in metazoa, such as *C. elegans* (0.01-0.4%), *Drosophila* (0.001-0.07%), *Xenopus laevis* (0.00009%), and mouse (0.0006-0.007%) (Greer et al., 2015; Koziol et al., 2015; Wu et al., 2016; Zhang et al., 2015), although it accumulates to a high level (0.1-0.2%) during vertebrate embryogenesis (Liu et al., 2016). Ciliates offer ideal systems for probing m^6^dA function, given the high abundance of m^6^dA and the availability of genetic and biochemical tools.

Intriguingly, in green algae and the ciliate *Tetrahymena*, m^6^dA is enriched in nucleosome linker regions (Fu et al., 2015; Hattman et al., 1978; Karrer and VanNuland, 1999; Pratt and Hattman, 1981, 1983; Wang et al., 2017), suggesting a role for m^6^dA in chromatin organization, or *vice versa*. Yet the functional relationship between m^6^dA and nucleosomes - if any - has remained unclear. Does m^6^dA directly disfavor nucleosomes, and if so, is it a graded or binary effect? Does the presence of histone post-translational modifications and ATP-dependent chromatin remodelers - both integral components of chromatin *in vivo* - modulate this interaction?

Here we address these questions by building synthetic, epigenetically defined chromosomes. Specifically, we generate chromosomes that either lack m^6^dA, or contain the modification at positions identical to their *in vivo* configuration. We also prepare chromosomes with *bona fide* telomeres that either lack or contain telomeric protein complexes. Using this library of synthetic chromosomes, we show that m^6^dA directly disfavors nucleosome occupancy in a quantitative, site-specific manner. Furthermore, this effect is modulated by histone post-translational modifications and chromatin remodelers, and is similar in magnitude to that imposed by telomeric protein complexes. Together, we demonstrate the utility of *Oxytricha* chromosomes as a versatile platform for functionally dissecting epigenetic modifications in native DNA.

## Results

### Epigenomic profiles of chromatin and transcription in *Oxytricha*

We generated genome-wide *in vivo* maps of nucleosome positioning, transcription, and m^6^dA in the *Oxytricha* macronuclear genome using MNase-seq, poly(A)+ RNA-seq, 5’-complete cDNA-seq, and single molecule real time (SMRT) sequencing (Figure 1). The presence of m^6^dA in *Oxytricha* DNA was independently validated by mass spectrometry (Figure S1). *Oxytricha* transcription start sites (TSSs) localize within 100bp of chromosome ends, upstream of a phased nucleosome array (Figure 1A). Strikingly, m^6^dA is enriched in three consecutive nucleosome depleted regions directly downstream of TSSs. Each cluster contains varying densities of m^6^dA (Figure 1B), with a maximum of 22 sites in the second cluster (Table S1A). Highly transcribed chromosomes tend to bear more m^6^dA, suggesting a positive role of this DNA modification in gene regulation (Figure 1C). Moreover, the majority of methylation was found on both DNA strands within an ApT motif (Figures 1D and 1E). m^6^dA occurs on sense and antisense strands with approximately equal frequency, indicating that the methylation machinery does not function strand-specifically.

**Figure 1.**
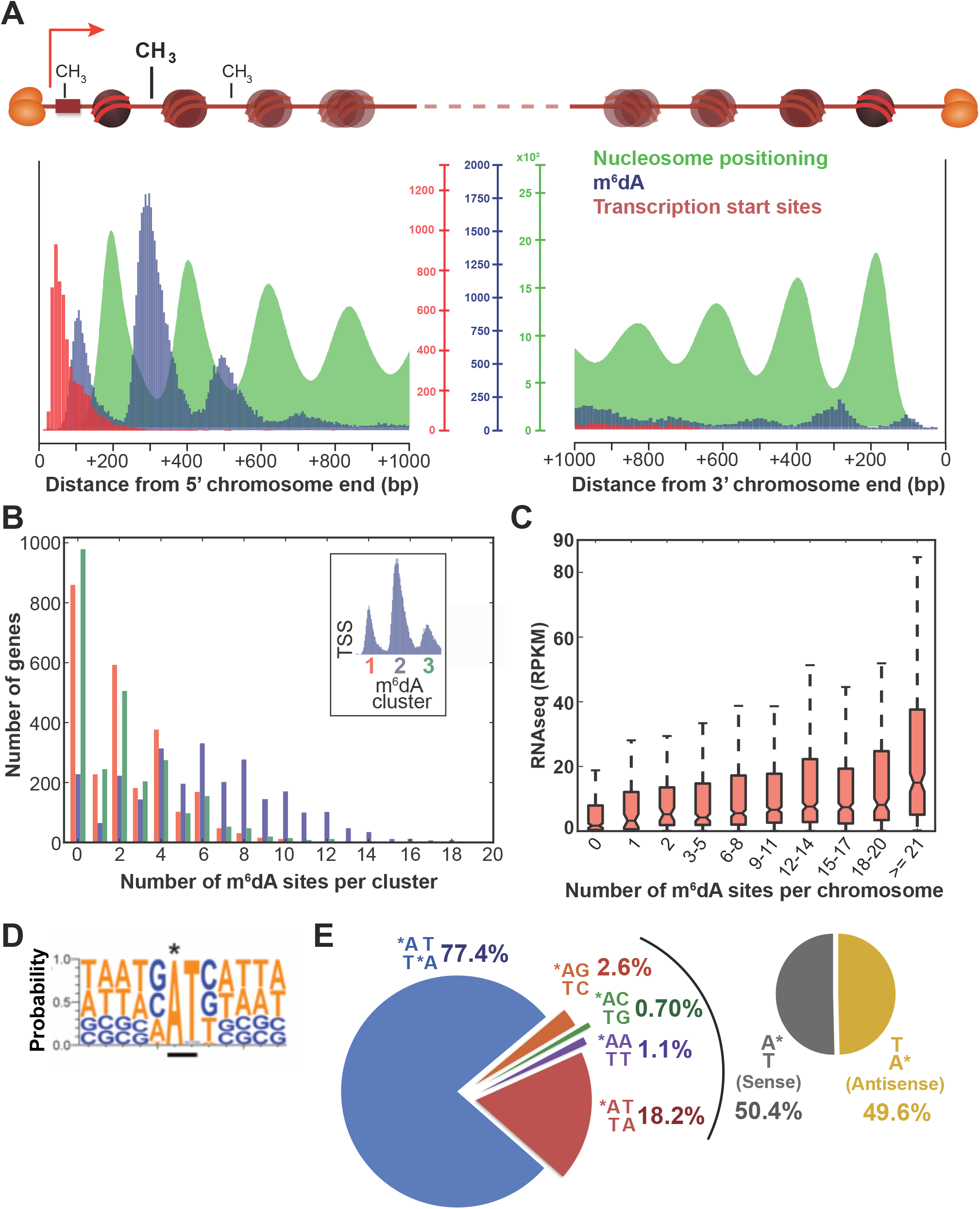
Epigenomic profiles of chromatin, transcription and DNA methylation in *Oxytricha* chromosomes. **(A)** Meta-chromosome plots overlaying MNase-seq (nucleosome occupancy *in vivo*), SMRT-seq (m^6^dA), and 5’-complete cDNA sequencing data (transcription start sites; TSSs) at *Oxytricha* chromosome ends. Heterodimeric telomere protein complexes protect each end *in vivo*, and are denoted as orange ovals. The 5’ end is designated as being proximal to TSSs. **(B)** Frequency of m^6^dA modifications downstream of individual TSSs. Histogram plots denote the distribution of m^6^dA frequencies within each aggregate cluster. **(C)** Transcriptional activity is positively correlated with the total number of m^6^dA within the corresponding chromosome. RNAseq data are derived from poly(A)-enriched RNA. RPKM denotes the number of reads per kilobase of chromosome per million mapped reads. **(D)** Composite analysis of 65,107 methylation sites reveals that m^6^dA occurs within an 5’-ApT-3’ dinucleotide motif. **(E)** Distribution of various m^6^dA dinucleotide motifs across the genome. Asterisk indicates DNA N6-methyladenine.

### m^6^dA localization in nucleosome linker regions of highly transcribed genes is deeply conserved across microbial eukaryotes

To better understand the conservation and evolutionary significance of m^6^dA, we also performed high coverage SMRT sequencing and mass spectrometry validation (Figures S1 and S2) in the ciliate *Tetrahymena*, which diverged from *Oxytricha* over 1 billion years ago (Bracht et al., 2013). The genome architecture of *Tetrahymena* is drastically different from *Oxytricha*, with chromosomes several orders of magnitude larger, spanning tens of kilobases to megabases in length (Coyne et al., 2008; Eisen et al., 2006). We find that m^6^dA is similarly enriched in nucleosome linker regions in *Tetrahymena*, consistent with earlier reports (Gorovsky et al., 1973; Hattman et al., 1978; Karrer and VanNuland, 1999; Pratt and Hattman, 1981). m^6^dA occurs in an ApT dinucleotide motif in both *Tetrahymena* and *Oxytricha* (Figures 1 and S2; also Bromberg et al., 1982), suggesting that the underlying enzymatic machinery responsible for m^6^dA deposition is conserved. Actively transcribed genes in *Tetrahymena* possess higher levels of m^6^dA, despite transcription start sites being distant from chromosome ends (Figure S2). m^6^dA is thus associated with transcribed DNA templates rather than proximity to telomeres *per se*. While m^6^dA patterns are broadly similar between *Oxytricha* and *Tetrahymena*, we note that its peak density is quantitatively different in *Tetrahymena* genes, and is considerably further downstream of TSSs than in *Oxytricha* (Figure 1A and Table S1A). Given that green algae possess a generally similar m^6^dA distribution and methylation motif as *Oxytricha* and *Tetrahymena* (Fu et al., 2015), we conclude that conserved mechanisms underlie m^6^dA establishment and function in ciliates and unicellular plants.

### m^6^dA directly disfavors nucleosome occupancy across the genome

Most strikingly conserved across the m^6^dA patterns of *Oxytricha, Tetrahymena,* and green algae is the inverse correlation between m^6^dA and nucleosome positioning *in vivo*. However, SMRT-seq data alone do not indicate causality. To test this directly, we exploited the naturally fragmented architecture of the *Oxytricha* macronuclear genome to amplify complete chromosomes using PCR. This erases all cognate m^6^dA, while fully preserving DNA sequence and physical linkage within each chromosome. We selected 98 unique chromosomes that collectively reflect overall genome properties, including AT content, chromosome length and transcriptional activity (Figure 2A; Table S1B). Only high copy number chromosomes were selected to ensure high-confidence identification of m^6^dA marks. Full-length chromosomes were individually PCR-amplified from genomic DNA, resulting in the collective erasure of 2,344 m^6^dA marks. Each chromosome was purified and subsequently mixed together in stoichiometric ratios to obtain a “minigenome” (Figure 2B). Native genomic DNA (containing m^6^dA) and amplified minigenome DNA (lacking m^6^dA) were each assembled into chromatin *in vitro* using *Xenopus* or *Oxytricha* histone octamers (Figures S3 and S4) and analyzed using MNase-seq. We computed nucleosome occupancy from the native genome and minigenome samples across 199,795 overlapping DNA windows, spanning all basepairs in the 98 chromosomes. This allowed the direct comparison of nucleosome occupancy in each window of identical DNA sequence, with and without m^6^dA (Figures 2C and 2D). Windows indeed exhibit lower nucleosome occupancy with increasing m^6^dA, confirming the quantitative nature of this effect. Furthermore, similar trends were observed for both native *Oxytricha* and recombinant *Xenopus* histones, suggesting that the effects of m^6^dA on nucleosome organization arise mainly from intrinsic features of the histone octamer rather than from species-specific variants (Figure 2C and 2D).

**Figure 2.**
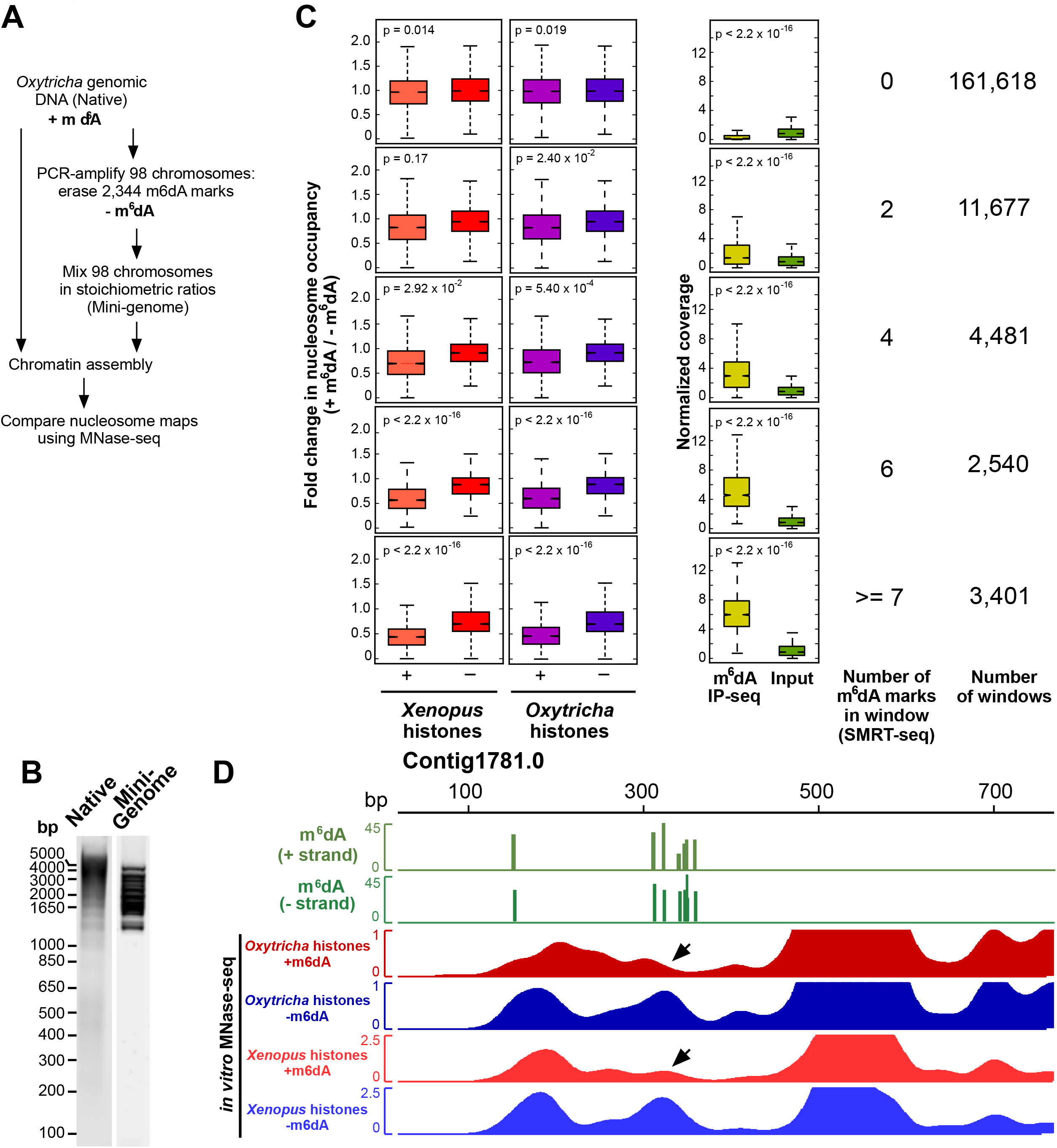
m^6^dA shapes nucleosome organization across the genome. **(A)** Experimental workflow. **(B)** Agarose gel analysis of *Oxytricha* gDNA (‘Native’) and mini-genome DNA before chromatin assembly. **(C)** Methylated regions of the genome exhibit lower nucleosome occupancy compared to identical DNA sequences lacking m^6^dA. Nucleosome occupancy *(in vitro* MNase-seq) and m^6^dA IP-seq coverage was calculated within overlapping 51bp windows across the 98 assayed chromosomes. Windows were binned according to the number of m^6^dA residues within. The MNase-seq coverage from chromatinized gDNA was divided by the corresponding coverage from chromatinized mini-genome DNA to obtain the fold change in nucleosome occupancy in each window (“+” histones). Naked gDNA and mini-genome DNA were also MNase-digested, sequenced and analyzed in the same manner to control for MNase sequence preferences (“-” histones). P values were calculated using a two-sample unequal variance t-test. **(D)** Tracks of m^6^dA distribution and MNase-seq coverage reveal a reduction in nucleosome occupancy at methylated loci (black arrowheads). “+” m^6^dA refers to chromatin assembled on *Oxytricha* gDNA, while “-” m^6^dA denotes chromatin assembled on mini-genome DNA. Similar trends are observed for both *Oxytricha* and *Xenopus* histones.

### Modular synthesis of an epigenetically defined chromosome

In principle, we reasoned that *Oxytricha* chromosomes could be constructed *de novo* via ligation of individual DNA building blocks, themselves generated in large quantities through PCR. The introduction of epigenetic modifications onto oligonucleotides before ligation would localize them to desired sites in the chromosome. We developed a streamlined chromosome synthesis scheme involving consecutive restriction enzyme digestion, ligation, and size selection steps (see Methods). Using this approach, we built a completely synthetic chromosome *in vitro* with a fully native DNA sequence, containing m^6^dA at all sites detected by SMRT-seq *in vivo*. We used this construct to dissect the effect of m^6^dA on nucleosome occupancy. The representative chromosome, *Contig1781.0*, is 1.3kb and contains a single highly transcribed gene with a clearly defined TSS (Figure 3A) - features characteristic of typical *Oxytricha* chromosomes. We independently validated the location of m^6^dA *in vivo* by sequencing chromosomal DNA immunoprecipitated with an anti-m^6^dA antibody (Figure 3A).

**Figure 3.**
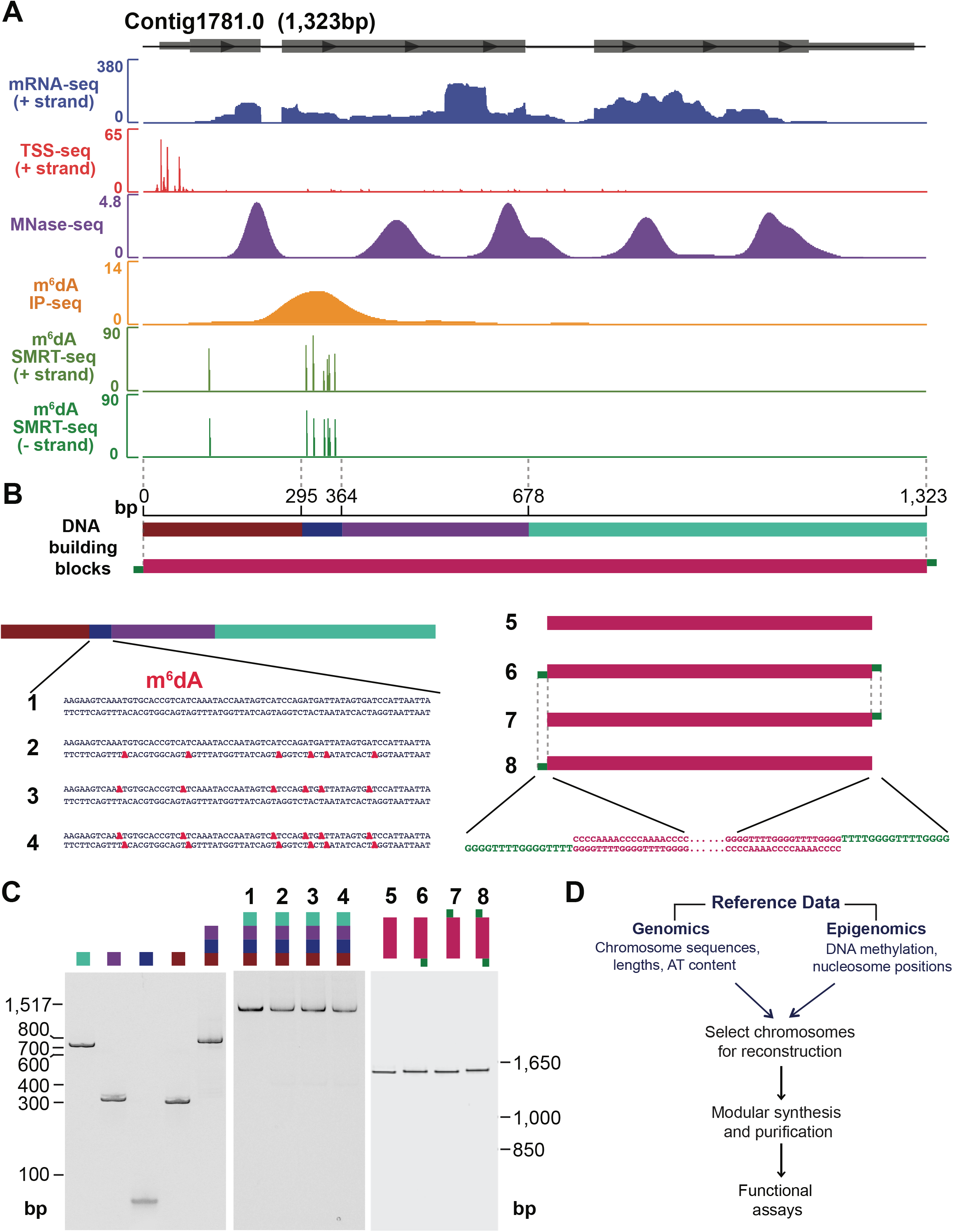
Modular synthesis of full-length *Oxytricha* chromosomes. **(A)** Chromatin and transcriptional features used to select, design and synthesize a full-length chromosome (Contig1781.0). Horizontal grey boxes represent the single annotated gene within this chromosome, with arrows denoting its orientation. All data tracks represent normalized sequencing coverage except for SMRT-seq, which represents the Phred-transformed p-value of detection of each methylated base. **(B)** Schematic of building blocks used to construct two sets of synthetic *Oxytricha* chromosomes. Different colors denote separate DNA building blocks ligated to form the 1,323bp full-length chromosome. Narrow green blocks in chromosomes 6 – 8 represent 16nt cognate single-stranded 3’ termini necessary for binding of the heterodimeric telomere protein complex. Each building block is drawn to scale. All m^6^dA residues lie in cognate positions discovered by SMRT-seq, and are highlighted with bold red. **(C)** Native polyacrylamide gel analysis of building blocks and purified synthetic chromosomes. **(D)** Workflow of chromosome design, synthesis, and application.

Four chromosome variants were synthesized, with cognate m^6^dA sites on neither, one, or both DNA strands (chromosomes 1 - 4 in Figures 3B, 3C, and S5A). Each chromosome was cloned and sequenced to verify the accuracy of construction (Figure S5C). With these chromosomal DNA templates in hand, we investigated the impact of m^6^dA on nucleosome occupancy. Chromatin was assembled by salt dialysis and subsequently digested with MNase to obtain mononucleosomal DNA (Figure 4A and S3). Tiling qPCR was used to quantify nucleosome occupancy at ~50bp increments along the entire length of the synthetic chromosome (Figure 4B). The fully methylated locus exhibits a ~46% reduction in nucleosome occupancy relative to the unmethylated variant, while hemimethylated chromosomes containing half the number of m^6^dA marks showed intermediate nucleosome occupancy at the corresponding region (Figures 4B and 4C). The reduction in nucleosome occupancy was confined to the methylated region, and not observed across the rest of the chromosome. We therefore conclude that m^6^dA directly disfavors nucleosome occupancy in a local, quantitative manner.

**Figure 4.**
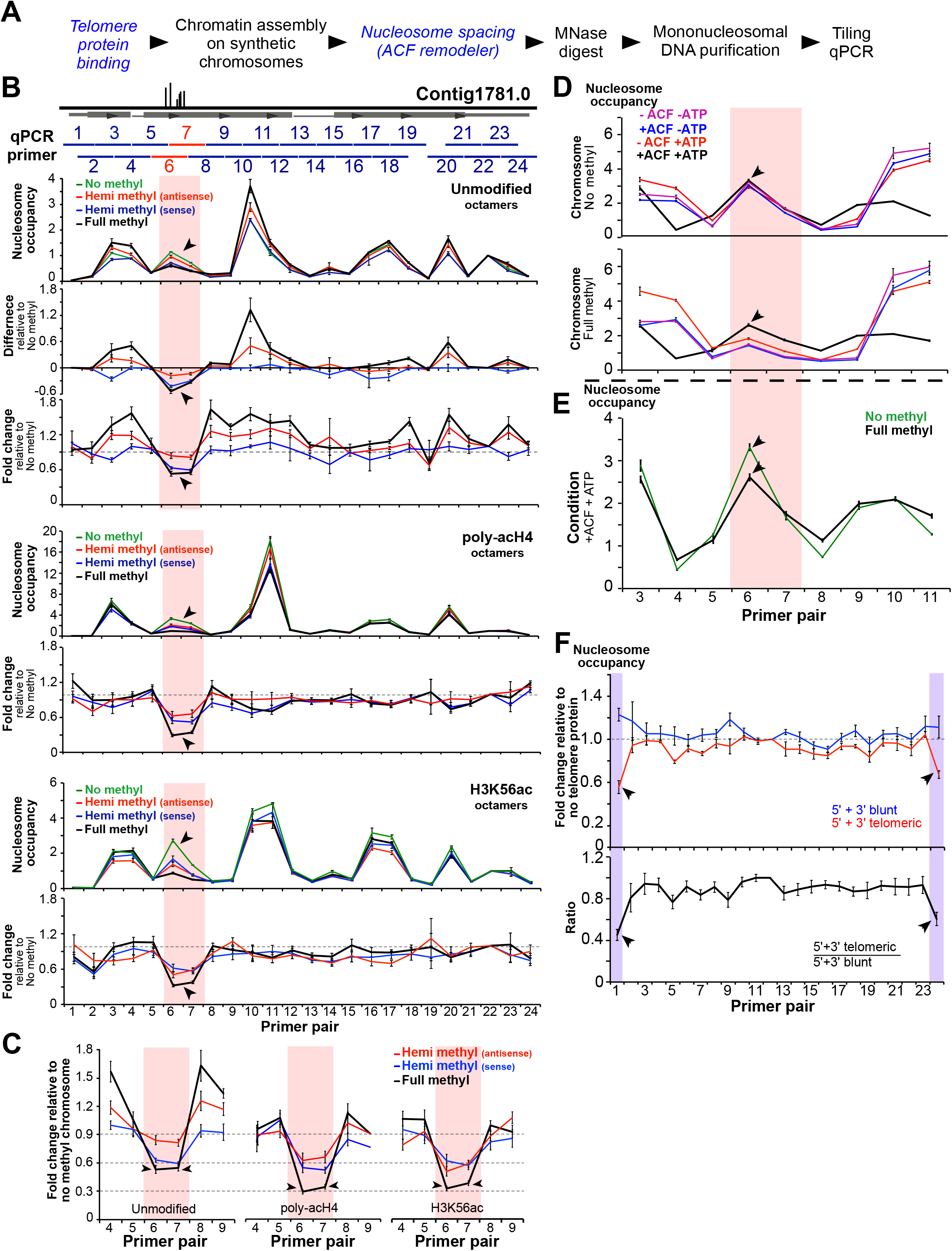
Quantitative modulation of nucleosome occupancy by m^6^dA in synthetic chromosomes. **(A)** Experimental workflow. Italicized blue steps are selectively included. **(B)** Tiling qPCR analysis of nucleosome occupancy in the synthetic chromosome Contig1781.0. Horizontal grey box represents the annotated gene within Contig1781.0. Horizontal blue bars span ~100bp regions amplified by qPCR primer pairs. Red horizontal lines and vertical bars represent the region containing m^6^dA. ‘Hemi methyl’ chromosomes contain m^6^dA on the antisense and sense strands respectively, while the ‘Full methyl’ chromosomes has m^6^dA on both strands. m^6^dA base positions are depicted in Figure 3B. Nucleosome occupancy represents normalized qPCR signal at each locus (see Methods). For each qPCR locus, “difference” = nucleosome occupancy in methylated chromosome - nucleosome occupancy in no methyl chromosome; “fold change” = nucleosome occupancy in methylated chromosome / nucleosome occupancy in no methyl chromosome. The decrease in nucleosome occupancy specifically at the m^6^dA cluster (black arrowheads) is observed for all octamer types. **(C)** The methylated region exhibits a greater decrease in nucleosome occupancy with H3K56 or poly-acH4 (black arrowheads) compared to unmodified octamers. **(D)** The chromatin remodeler ACF can partially overcome the effects of m^6^dA on nucleosome organization in an ATP-dependent manner. ACF equalizes nucleosome occupancy between the m^6^dA cluster and flanking regions in the presence of ATP (black line), resulting in a relative increase in nucleosome occupancy over the methylated region. **(E)** Nucleosome occupancy at the methylated locus is not restored to the same level as the unmethylated control (black arrowheads) in the presence of ACF and ATP. **(F)** Telomere binding proteins (TeBPs) disfavor nucleosome occupancy at chromosome ends (black arrowheads). The 5’+3’ telomeric chromosome possesses terminal 16nt single-stranded telomeric sequence necessary for TeBP binding, while the 5’+3’ blunt chromosome lacks such tails. For each chromosome, nucleosome occupancy in the presence of TeBPs was normalized relative to the absence of TeBPs (see Methods). Error bars in all panels represent s.e.m. (n = 3-4).

### Chromatin remodelers partially restore nucleosome occupancy over m^6^dA sites

Nucleosome occupancy *in vivo* is influenced not only by DNA sequences, but also by *trans*-acting factors. ATP-dependent chromatin remodeling factors modulate nucleosome organization and help establish canonical nucleosome patterns near TSSs (Struhl and Segal, 2013). We used our synthetic methylated chromosomes to test how the well-studied chromatin remodeler ACF responds to m^6^dA in native DNA. ACF generates regularly spaced nucleosome arrays *in vitro* and *in vivo* (Clapier and Cairns, 2009; Ito et al., 1997). Its catalytic subunit ISWI is conserved across eukaryotes, including the ciliates *Oxytricha* and *Tetrahymena* (Figure S6). Synthetic chromosomes were assembled into chromatin by salt dialysis as before, then incubated with ACF in the presence of ATP (Figure S3D). We find that ACF partially - but not completely - restores nucleosome occupancy over the methylated locus in an ATP-dependent manner (Figure 4D and 4E). Chromatin remodelers may thus modulate nucleosome occupancy in concert with m^6^dA, each imposing distinct but interrelated effects.

### Histone acetylation and m^6^dA exert synergistic effects on nucleosome occupancy

Nucleosomes are decorated with post-translational modifications (PTMs) *in vivo*, which collectively modulate chromatin structure and function. A well-documented example is lysine acetylation, particularly at histone H3 lysine 56 (H3K56ac) and at multiple residues in the histone H4 N-terminal tail (poly-acH4). H3K56 lies at the entry and exit sites of a nucleosome, and its acetylation lowers the affinity of H3-H4 tetramers for DNA (Andrews et al., 2010) and increases DNA unwrapping (Simon et al., 2011), together leading to nucleosome destabilization. H4 polyacetylation reduces the net positive charge of histone octamers, weakening histone-DNA contacts (Hong et al., 1993). It also hinders chromatin compaction and nucleosome aggregation, indicating a role in regulating higher order chromatin structure (Allahverdi et al., 2011).

Since m^6^dA is embedded within chromatin *in vivo* and likely occurs in the context of histone PTMs, we asked whether the effects of m^6^dA on nucleosome positioning are themselves modulated by PTMs such as H3K56ac and poly-acH4, which influence chromatin structure. Using quantitative mass spectrometry, we verified that *Oxytricha* histones contain H3K56ac and poly-acH4 *in vivo*, along with numerous other sites of acetylation on H3 and H4 (Figure S7). We prepared recombinant H3K56ac and semisynthetic poly-AcH4 using amber codon suppression and native chemical ligation, respectively (Figure S4). The modified histones were refolded into octamers in parallel with unmodified controls and subsequently assembled into chromatin on the methylated chromosomes. Different histone octamer types exhibited qualitatively similar patterns of nucleosome organization across each chromosome (Figure 4B). Curiously, poly-acH4-containing octamers gave rise to higher nucleosome occupancy near the center of the chromosome relative to flanking regions, despite a weaker affinity of these octamers for DNA *per se*. Since the central region is intrinsically favorable for nucleosome formation (occupancy is highest in this region, even for unmodified octamers), it may be less sensitive to decreases in octamer affinity compared to flanking regions. We then computed the fold-change in nucleosome occupancy in methylated chromosomes relative to the unmethylated control of identical DNA sequence. This calculation was performed separately for each octamer type. Chromatin assembled with H3K56ac or poly-acH4 exhibited a significantly greater reduction in nucleosome occupancy than unmodified octamers, for fully methylated DNA (Figure 4C). Therefore, H3K56ac and poly-acH4 can act synergistically with m^6^dA to disfavor nucleosome occupancy. These data broadly reflect the complex interplay between histone PTMs and m^6^dA in modulating chromatin structure. In our model, H3K56ac and poly-acH4 may not actually localize near m^6^dA within this specific chromosome *per se*, but we propose that histone PTMs that alter chromatin structure can work in concert with m^6^dA to modulate nucleosome organization.

### Telomere proteins and m^6^dA disfavor nucleosome occupancy to similar extents

The synthetic chromosomes described thus far contain blunt telomeric ends, but *Oxytricha* chromosome termini *in vivo* possess 16nt single-stranded 3’ DNA tails, necessary for associating with the telomere end-binding protein complex, TeBPα and TeBPβ, together similar in mass to a histone octamer. TeBPα/β bind cooperatively to the single-stranded telomeric tail to form a ternary complex (K_d_ = 2nM^2^), stable even in *2M* NaCl (Gottschling and Zakian, 1986; Horvath et al., 1998). To determine whether telomere protein binding at chromosome termini influences nucleosome occupancy, and to compare its effects to m^6^dA, we used our modular synthesis scheme to build synthetic chromosomes with *bona fide* 3’ tails (chromosomes 6 - 8 in Figure 3B, 3C and S5B). Recombinantly expressed and purified *Oxytricha* TeBPα and TeBPβ (Figure 5A) were both shown, in methylation protection assays, to bind cooperatively to *Oxytricha* gDNA termini, yielding guanine residue protection patterns consistent with previous studies (Gray et al., 1991) (Figure 5C). TeBPα and TeBPβ were then pre-bound to synthetic chromosomes and subsequently used for chromatin assembly via salt dialysis (Figure 5D). Gel shift assays confirmed that both subunits remain associated with synthetic chromosome ends in 2M NaCl, the highest salt concentration in the chromatin assembly procedure (Figures 5E and 5F). We also verified that telomere protein binding occurs independently at 5’ and 3’ chromosome termini (Figures 5E and 5F). The synthetic chromosome pre-bound with TeBP proteins at both termini exhibits a 40-50% decrease in nucleosome occupancy, within 50bp of each chromosome end (Figure 5F). This region directly overlaps with 32.1% of transcription start sites in the genome (Figure 1A), and may thus influence promoter chromatin accessibility. Indeed, TeBP binding has been reported to modulate transcription initiation in *Euplotes crassus* (Bender and Klein, 1997), a ciliate with similar chromosome architecture to *Oxytricha*. The observed reduction in nucleosome occupancy upon TeBP binding is quantitatively similar to that imposed by fully methylated chromosomal loci (Figure 4F). Therefore, both telomere protein binding and m^6^dA deposition sculpt nucleosome organization, though m^6^dA exerts a graded rather than all-or-none effect, depending on the number of m^6^dA marks present.

**Figure 5.**
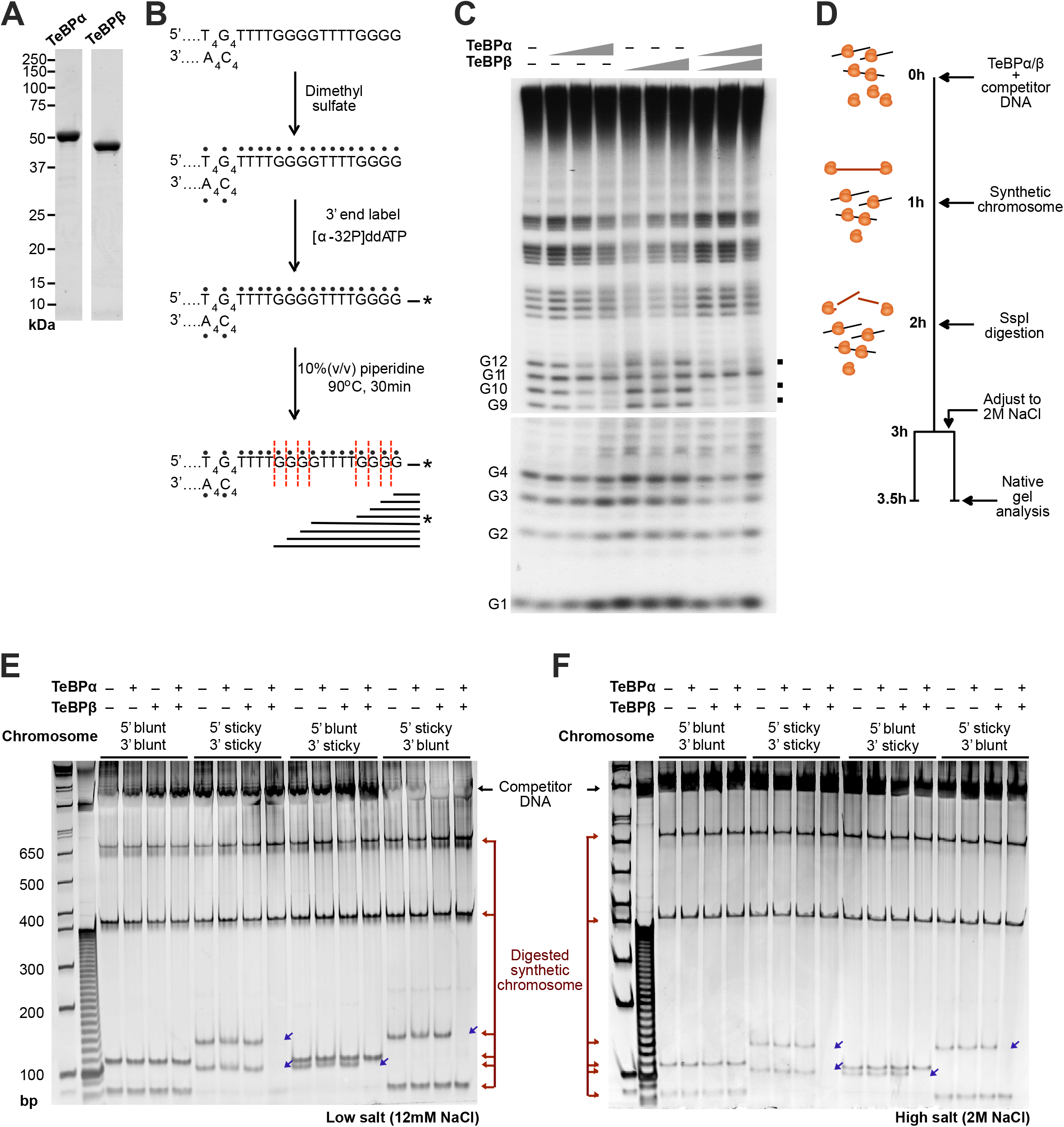
Preparation of synthetic *Oxytricha* chromosomes bound with telomere proteins. **(A)** SDS-PAGE analysis of recombinant *Oxytricha* TeBPα and TeBPβ proteins. **(B)** Methylation protection assay to test TeBPα/β activity. The 3’ ends of *Oxytricha* genomic DNA bear 16nt telomeric tails, as illustrated. The DNA is treated with dimethyl sulfate, which methylates individual DNA bases (represented by black dots). Methylated DNA is subsequently 3’ end-labeled and heated in piperidine, resulting in cleavage specifically at methylated guanine residues (vertical red lines). The cleavage products are then resolved by 20% urea-PAGE, giving rise to a “ladder” of DNA fragments that can be visualized by autoradiography. **(C)** Cooperative TeBPα/β binding to *Oxytricha* genomic DNA before dimethyl sulfate treatment protects specific guanine residues from methylation and thus piperidine cleavage (marked with black squares). Methylation protection is observed at guanines G9, G10, and G12 in the presence of both TeBPα and TeBPβ, consistent with previous reports. TeBPα is known to exhibit similar but weaker DNA-binding effects in the absence of TeBPβ. **(D)** Telomere proteins are pre-incubated with non-specific competitor DNA before adding synthetic chromosomes that bear single-stranded telomeric tails. This results in specific binding of TeBPα/β to synthetic chromosome ends. These DNA-protein complexes can then be directly used for chromatin assembly. Alternatively, to verify TeBPα/β binding, each sample is treated with a restriction enzyme that cleaves near synthetic chromosome ends. Liberated DNA fragments are visualized by native PAGE. **(E)** Synthetic chromosome ends that bear the 16 nt telomeric tail (“sticky”) are supershifted into the well in the presence of both TeBPα/β, while those lacking the tail (“blunt”) are unaffected. Blue arrows indicate supershifted fragments. Binding conditions are in 12 mM NaCl. **(F)** Similar results are observed in 2 M NaCl, consistent with previous reports that TeBPα/β binding is unaffected by high salt concentrations.

## Discussion

We report that m^6^dA directly disfavors nucleosome occupancy and that this effect can be modulated by histone post-translational modifications and ATP-dependent chromatin remodelers. We expect the biochemical impact of m^6^dA to be directly pertinent across a wide diversity of eukaryotic genomes, including vertebrates, *C. elegans, Drosophila* and fungi, where this epigenetic modification has recently been documented (Greer et al., 2015; Koziol et al., 2015; Liu et al., 2016; Mondo et al., 2017; Wu et al., 2016; Zhang et al., 2015). The current experiments do not reveal exactly how m^6^dA disfavors nucleosome occupancy. Early studies suggest that N6-methylation destabilizes dA:dT base pairing, leading to a decrease in the melting temperature of DNA (Engel and von Hippel, 1978). Whether this or some other physico-chemical property of m^6^dA contributes to lowered nucleosome stability awaits further investigation.

While m^6^dA directly disfavors nucleosome occupancy, it may also be possible that nucleosomes in turn inhibit m^6^dA deposition by the putative methylase, establishing a positive feedback loop that reinforces the inverse relationship between nucleosome occupancy and DNA methylation. Aside from simple physical accessibility, the activity of the m^6^dA methylase may differ upon binding to nucleosomes, compared to naked DNA. Histone variants and PTMs commonly enriched near transcription start sites may also modulate the enzyme’s activity. Future identification of the ciliate m^6^dA methylase would shed light on these questions and advance our understanding of how nucleosomes and DNA methylation interact to establish chromatin architecture near TSSs.

What could be the identity of the m^6^dA methylase in ciliates? While typical Dam and DMNT-like DNA methylases are absent from the *Oxytricha* and *Tetrahymena* genomes, there are 5-6 candidate genes with predicted MT-A70 domains, homologous to the METTL gene family (Iyer et al., 2016; Luo et al., 2015). Although some mammalian MT-A70 proteins are known to catalyze m^6^A methylation on RNA, they may have been co-opted to deposit methyl groups on DNA substrates in ciliates. Functional perturbations of these candidates *in vivo* would test such predictions. Equally intriguing is the observation that actively transcribed genes possess high levels of m^6^dA. This trend is deeply conserved, being present in the distantly related ciliates *Tetrahymena* and *Oxytricha*, as well as green algae and basal fungi. It is possible that the m^6^dA methylase is associated with RNA polymerases, resulting in m^6^dA deposition during transcriptional elongation. Alternatively, transcription factors may contain “reader” domains that specifically recognize m^6^dA, thus increasing transcriptional output at methylated loci. Importantly, these two scenarios are not mutually exclusive. We envision that the use of synthetic *Oxytricha* chromosomes, in conjunction with transcriptionally competent nuclear extracts, would constitute an especially useful biochemical tool for dissecting such effects.

More broadly, our study showcases the utility of *Oxytricha* chromosomes in advancing chromatin biology. Each chromosome essentially comprises a nucleosome array, capped with telomeric protein complexes at both ends. Here we show how these features can be reconstructed in their entirety using synthetic chromosomes. By extending current technologies (Müller et al., 2016), it should be feasible to introduce both modified nucleosomes and DNA methylation in a site-specific manner on full-length chromosomes. Such ‘designer’ constructs will serve as powerful tools for studying DNA-templated processes such as transcription within the context of a fully native DNA environment.

## Author contributions

L.Y.B. conceived the project, designed research, synthesized chromosomes, performed computational and experimental analysis for all Figures and Tables, and wrote the manuscript. G.T.D. designed research, synthesized chromosomes, and prepared all *Xenopus* histones. K.A.L. processed raw SMRT-seq data. K.K. and B.A.G performed mass spectrometry analysis of *Oxytricha* histones. E.R.H. performed m^6^dA IP-seq. J.R.B. prepared *Oxytricha* DNA for SMRT-seq. R.P.S performed SMRT-seq. T.W.M. and L.F.L. conceived the project, designed research and analyzed data. G.T.D, T.W.M, and L.F.L. edited the manuscript.

## Acknowledgments

We thank Geoffrey Dann for discussions regarding the ACF remodeler and for performing preliminary tests of its activity; Tharan Srikumar for assistance with nucleoside mass spectrometry; Wei Wang, Jessica Wiggins, and Jennifer Miller for assistance with Illumina sequencing; C. David Allis and Virginia Zakian for suggestions and comments on the project; Ana Mostafavi and Glen Liszczak for advice regarding amber codon suppression, and Jingmei Wang, Barbara Dul, and Fei Song for general laboratory support. This work was funded by NIH grants R01-GM59708 to L.F.L and R01-GM107047 to T.W.M.

---

**Table S1. Descriptive statistics**

**(A)** Properties of m^6^dA distribution near TSSs. An m^6^dA site is classified as lying within a particular methyl cluster if it is within 50 bp of the peak derived from the aggregate m^6^dA distribution. Aggregate m^6^dA peak positions in *Oxytricha* are +56 bp, +235 bp, and +436 bp downstream of the TSS, while those in *Tetrahymena* are +205 bp, +400 bp, and +597 bp respectively.

**(B)** Properties of *Oxytricha* chromosomes in native genomic DNA and mini-genome DNA.

**Table S2. Primer sequences**

All primers are in the 5’ to 3’ direction.

**Figure S1. Mass spectrometry analysis confirms the presence of m^6^dA in ciliate DNA**

*Oxytricha* and *Tetrahymena* genomic DNA were digested into nucleosides and used for reverse-phase HPLC and subsequent mass spectrometry. Chemically synthesized dA and m^6^dA were used as standards, with 12 pmol and 0.3 pmol respectively loaded. Eluted peaks with expected masses of m6dA are detected in both *Tetrahymena* and *Oxytricha* nucleosides.

**Figure S2. Genomic distribution of m^6^dA in *Tetrahymena thermophila***

MNase-seq data and RNA-seq data were obtained from previously published datasets (Beh et al., 2015; Xiong et al., 2012).

**(A)** Meta-chromosome plots overlaying MNase-seq (nucleosome positioning in vivo) and SMRT-seq (m^6^dA), relative to transcription start sites. m^6^dA lies mainly within nucleosome linker regions, between the +1, +2, and +3 nucleosomes.

**(B)** Frequency of m^6^dA modifications downstream of TSSs.

**(C)** Transcriptional activity is positively correlated with the total number of m^6^dA within the corresponding gene. RPKM denotes the number of reads per kilobase per million mapped reads.

**(D)** Composite analysis of 441,618 methylation sites reveals that m^6^dA occurs within an 5’-ApT-3’ dinucleotide motif in *Tetrahymena*, consistent with isotopic labeling experiments (Bromberg et al., 1982; Wang et al., 2017) and similar to *Oxytricha*.

**(E)** Distribution of various m^6^dA dinucleotide motifs across the genome.

**(F)** Organization of transcription, nucleosome organization, and m^6^dA in a single *Tetrahymena* gene.

**Figure S3. Gel analysis of assembled chromatin**

*Xenopus* or *Oxytricha* histone octamers were assembled on DNA through salt dialysis and subsequently digested with MNase to obtain monunucleosome-sized fragments. The resulting products were analyzed by agarose gel electrophoresis.

**(A)** Chromatin assembled with synthetic chromosome DNA. All assemblies were performed in the presence of an approximately 100-fold mass excess of buffer DNA relative to synthetic chromosome (see Methods). Representative assemblies with the unmethylated chromosome are shown. Methylated chromosome assemblies were separately performed in place of the unmethylated variant.

**(B)** Chromatin assembled on PCR-amplified mini-genome DNA.

**(C)** Chromatin assembled on native genomic DNA.

**(D)** Chromatin assembled on unmethylated synthetic chromosomes and incubated with ACF and ATP. Regularly spaced nucleosomes are observed only when ACF and ATP are present.

**Figure S4. Preparation of histone octamers for chromatin assembly**

*Xenopus* unmodified core histones were expressed recombinantly, while H3K56ac and poly-acH4 were synthesized through amber codon suppression and native chemical ligation, respectively. *Oxytricha* histones were acid-extracted from vegetative nuclei. *Oxytricha* and *Xenopus* histones were subsequently refolded into octamers and purified through size exclusion chromatography.

**(A)** Reverse-phase HPLC analysis of purified *Xenopus* poly-acH4.

**(B)** ESI mass spectrometry analysis of purified *Xenopus* poly-acH4.

**(C)** Reverse-phase HPLC analysis of purified *Xenopus* H3K56ac.

**(D)** ESI mass spectrometry analysis of purified *Xenopus* H3K56ac.

**(E)** Reverse-phase HPLC purification of acid-extracted *Oxytricha* histones. Fractions 1-5 were individually collected and analyzed by Coomassie staining and western blotting.

**(F)** SDS-PAGE analysis of purified *Oxytricha* histone fractions.

**(G)** Western blot analysis confirms identity of each *Oxytricha* histone fraction.

**(H)** SDS-PAGE analysis of purified histone octamers.

**Figure S5. Schematic of chromosome synthesis strategy**

Staggered dotted lines represent BsaI cleavage sites.

**(A)** Assembly of methylated chromosome variants.

**(B)** Assembly of chromosome variants with 3’ single-stranded telomeric tails.

**(C)** Sanger sequencing of cloned synthetic chromosomes. Continuous horizontal green bar represents full sequence identity between the reference chromosome and individual sequencing reads.

**Figure S6. Putative ciliate ISWI orthologs**

ISWI is a member of the SW12/SNF2 ATPase family that acts as chromatin remodelers. The *Oxytricha* and *Tetrahymena* genomes were queried by BLASTP using *Drosophila melanogaster* ISWI (UniProt ID: Q24368), and the reciprocal best hit was retrieved from each genome. BLASTP e-values were 0.0 in each case. Putative *Tetrahymena* and *Oxytricha* orthologs were queried for protein domains and associated GO terms using InterPro (Finn et al., 2017). ISWI contains an N-terminal catalytic ATPase domain, and a C-terminal HAND-SANT-SLIDE module necessary for nucleosome binding and mobilization.

**Figure S7. Detection of poly-acH4 and H3K56ac in *Oxytricha* cells using quantitative mass spectrometry**

**(A)** Middle-down mass spectrometry (MS) quantification of histone H3 variants. Histone H3 variants are listed along the y-axis, and are henceforth abbreviated as g60, g122, g137, g54, g33, and g10 respectively. x-axis represents total ion count (arbitrary units).

**(B)** Middle-down MS quantification of histone H3 (Contig4701.0.g33) acetylation. Data from four biological replicates are respectively shown. Positions of PTMs are listed along the *x*-axis. *y*-axis represents the cumulative abundance of acetylation on the modified Lys-residues as relative to the total histone H3. Each bar represents the averaged relative abundance (%) of 3 technical replicates (with exception of n_t_Rep4_=2); error bars represent ± standard deviation (stdev) of technical (n_t_Rep1-3_ = 3; n_t_Rep4_ = 2;) replicates.

**(C)** Bottom-up MS quantification of histone H3 acetylation. Positions of PTMs are listed along the *x*-axis. *y*-axis represents the cumulative abundance of acetylation on each residue. Histone peptides containing H3K56ac are KYQKSTELLIR (g122); KFQKSTELLIR (g10); KYQKSTDLLIR (g60); and RFQKSTELLIR (g33, g54, g137). Each bar represents the averaged relative abundance (%) of 4 biological replicates; n_b_ = 4). Error bars represent ± standard deviation (stdev) of biological replicates (n_b_ = 4).

**(E)** Bottom-up MS quantification of histone H4 acetylation. Positions of PTMs are listed along the *x*-axis. *y*-axis represents the cumulative abundance of acetylation on the four modified residues of the H4 peptide GKVGKGYGKVGAKR. The *Oxytricha* genome contains two annotated histone H4 genes with identical amino acid sequence.

**(F)** modified peptides from (d) as relative to total histone H4. Each bar represents the averaged relative abundance (%) of 4 biological replicates; n_b_ = 4). Error bars represent ± standard deviation (stdev) of biological (n_b_ = 4) replicates.

**Figure S8. Tiling qPCR analysis of nucleosome occupancy in spike-in and homogeneous synthetic chromosome preparations**

The blunt, unmethylated synthetic chromosome (construct #1 in Figure 3B) was used for chromatin assembly with (“Spike-in”) or without (“Homogeneous”) a hundred-fold excess of carrier DNA. In the latter case, an equivalent mass of synthetic chromosome was added in place of carrier DNA to maintain the same DNA concentration for chromatin assembly. Red and purple regions depict the corresponding regions where m^6^dA and TeBPα/β respectively modulate nucleosome occupancy in separate synthetic chromosomes studied in Figure 4. Black arrowheads indicate no decrease in nucleosome occupancy in these regions when carrier DNA is used.

## STAR Methods

### Contact for reagent and resource sharing

Further information and requests for resources and reagents should be directed to and will be fulfilled by the Lead Contact, Laura Landweber

### Experimental model and subject details

#### Oxytricha trifallax

Vegetative *Oxytricha trifallax* strain JRB310 was cultured at a density of 1.5 × 10^7^ cells/L to 2.5 × 10^7^ cells/L in Pringsheim media (0.11 mM Na_2_HPO_4_, 0.08 mM MgSO_4_, 0.85 mM Ca(NO_3_)_2_, 0.35 mM KCl, pH 7.0) and fed daily with *Chlamydomonas reinhardtii*. Cells were filtered through cheesecloth to remove debris and collected on a 10 μm Nitex mesh for subsequent experiments.

#### Tetrahymena thermophila

Vegetative *Tetrahymena thermophila* strain SB210 was cultured in 1xSPP media as previously described (Beh et al., 2015) at a density of ~35x10^4^ cells/mL and collected by centrifuged at 1000 x g for 5 min for subsequent experiments.

### Method details

#### *in vivo* MNase-seq

3 x 10^5^ vegetative *Oxytricha* cells were fixed in 1% (w/v) formaldehyde for 10 min at room temperature with gentle shaking, and then quenched with 125 mM glycine. Macronuclei were subsequently isolated by sucrose gradient centrifugation from fixed cells as previously described (Lauth et al., 1976). Purified nuclei were pelleted by centrifugation at 4000 x g, washed in 50 ml TMS buffer (10 mM Tris pH 7.5, 10 mM MgCl_2_, 3 mM CaCl_2_, 0.25 *M* sucrose), resuspended in a final volume of 300 μL, and equilibriated at 37°C for 5 min. Chromatin was then digested with MNase (New England BioLabs) at a final concentration of 15.7 Kunitz Units / μL at 37°C for 1 min 15 sec, 3 min, 5 min, 7 min 30sec, 10 min 30 sec, and 15 min respectively. Reactions were stopped by adding 1/2 volume of PK buffer (300 mM NaCl, 30 mM Tris pH 8, 75 mM EDTA pH 8, 1.5% (w/v) SDS, 0.5 mg/mL Proteinase K). Each sample was incubated at 65°C overnight to reverse crosslinks and deproteinate samples. Subsequently, nucleosomal DNA was purified through phenol:chloroform:isoamyl alcohol extraction and ethanol precipitation. Each sample was loaded on a 2% agarose-TAE gel to check the extent of MNase digestion. The sample exhibiting ~80% mononucleosomal species was selected for MNase-seq. analysis, in accordance with previous guidelines (Zhang and Pugh, 2011). Mononucleosome-sized DNA was gel-purified using a QIAquick gel extraction kit (QIAGEN) for subsequent paired-end sequencing on an Illumina HiSeq 2500 according to manufacturer’s instructions. All *Tetrahymena* MNase-seq data were obtained from (Beh et al., 2015).

### poly(A)+ RNA-seq and 5’-complete cDNA-seq

Vegetative *Oxytricha* cells were lysed in TRIzol reagent (Thermo Fisher Scientific) for total RNA isolation according to manufacturer’s instructions. Poly(A)+ RNA was then purified using the NEBNext Poly(A) mRNA Magnetic Isolation Module (New England BioLabs). The *Oxytricha* poly(A)+ RNA was prepared for RNA-seq using the ScriptSeq v2 RNA-Seq Library Preparation Kit (Illumina). *Tetrahymena* poly(A)+ RNA-seq data was obtained from (Xiong et al., 2012).

The 5’ ends of capped RNAs were enriched from vegetative *Oxytricha* total RNA using the RAMPAGE protocol (Batut et al., 2013). They were used for library preparation, Illumina sequencing and subsequent transcription start site determination (ie. “TSS-seq”). *Tetrahymena* transcription start site positions were obtained from (Beh et al., 2015).

### m^6^dA IP-seq

Genomic DNA was isolated from vegetative *Oxytricha* cells using the Nucleospin Tissue Kit (Macherey-Nagel). DNA was sheared into 150 bp fragments using a Covaris LE220 ultra-sonicator (Covaris). Samples were gel-purified on a 2% agarose-TAE gel, blunted with Klenow polymerase (New England Biolabs, MA), and purified using MinElute spin columns (QIAGEN). The fragmented DNA was dA-tailed using Klenow Fragment (3’ -> 5’ exo-) (New England BioLabs) and ligated to Illumina adaptors following manufacturer’s instructions. Subsequently, 2.2μg of adaptor-ligated DNA containing m^6^dA was immunoprecipitated using an anti-N6-methyladenosine antibody (m^6^A; Synaptic Systems) conjugated to Dynabeads (Life Technologies). The anti-m^6^A antibody is commonly used for RNA applications, but has also been demonstrated to recognize m^6^dA in DNA (Fioravanti et al., 2013; Xiao and Moore, 2011). The immunoprecipitated and input libraries were respectively treated with proteinase K, phenol:chloroform extracted, and ethanol precipitated. Finally, they were PCR-amplified using Phusion Hot Start polymerase (New England BioLabs) and used for Illumina sequencing.

### Sample preparation for SMRT-seq

Macronuclei were isolated from vegetative *Oxytricha* and *Tetrahymena* as previously described (Lauth et al., 1976) and used for genomic DNA isolation with the Nucleospin Tissue Kit (Macherey-Nagel). SMRT-seq was performed as previously described (Chen et al., 2014), according to manufacturer’s instructions, using P5-C3 and P6-C4 chemistry.

### Illumina data processing

Reads from all biological replicates from each dataset were merged before downstream processing. All Illumina sequencing data were quality trimmed (minimum quality score = 20) and length-filtered (minimum read length = 40 nt) using Galaxy (Blankenberg et al., 2010; Giardine et al., 2005; Goecks et al., 2010). MNase-seq and m^6^dA IP-seq reads were mapped to complete chromosomes in the *Oxytricha trifallax* JRB310 (August 2013 build) or *Tetrahymena* thermophila SB210 macronuclear reference genomes (June 2014 build) using Bowtie 2 (Langmead and Salzberg, 2012) with default settings, while poly(A)+ RNA-seq and TSS-seq reads were mapped using TopHat2 (Mortazavi et al., 2008) with August 2013 *Oxytricha* gene models or June 2014 *Tetrahymena* gene models, with default settings.

MNase-seq read pairs of length 122-172 bp were used for analysis. m^6^dA IP-seq single-end reads were extended to the mean fragment size, computed using cross-correlation analysis (Kharchenko et al., 2008). The per-basepair coverage of *Oxytricha* MNase-seq read pair centers (termed “nucleosome occupancy” in this manuscript) and full extended m^6^dA IP-seq reads were respectively computed across the genome. Subsequently, the per-basepair coverage values were normalized by the average coverage within each chromosome to account for differences in DNA copy number (and hence, read depth) between *Oxytricha* chromosomes (Swart et al., 2013). The per-bp coverage values were then smoothed using a Gaussian filter of standard deviation = 15 (Beh et al., 2015; Kaplan et al., 2010). For RNA-seq data, the number of reads per kilobase of chromosome per million mapped reads (RPKM) was calculated for each chromosome without normalization by DNA copy number since there is no correlation between *Oxytricha* DNA and transcript levels (Swart et al., 2013). *Oxytricha* TSS-seq data were processed using CAGEr (Haberle et al., 2015); with clusterCTSS parameters (threshold = 1.6, thresholdIsTpm = TRUE, nrPassThreshold = 1, method = “paraclu”, removeSingletons = TRUE, keepSingletonsAbove = 5). Only TSSs with tags per million counts > 0.1 were used for downstream analysis. Mapped *Tetrahymena* MNase-seq and RNA-seq data were processed as previously described (Beh et al., 2015).

### SMRT-seq data processing

We processed SMRT-seq data with SMRTPipe v1.87.139483 in the SMRT Analysis 2.3.0 environment using, in order, the P_Fetch, P_Filter (with minLength = 50, minSubreadLength = 50, readScore = 0.75, and artifact = -1000), P_FilterReports, P_Mapping (with gff2Bed = True, pulsemetrics = DeletionQV, IPD, InsertionQV, PulseWidth, QualityValue, MergeQV, SubstitutionQV, DeletionTag, and loadPulseOpts = byread), P_MappingReports, P_GenomicConsensus (with algorithm = quiver, outputConsensus = True, and enableMapQVFilter = True), P_ConsensusReports, and P_ModificationDetection (with identifyModifcations = True, enableMapQVFilter = False, and mapQvThreshold = 10) modules. All other parameters were set to the default. The *Oxytricha* August 2013 reference genome build was used for mapping Oxytricha SMRT-seq reads, with Contig10040.0.1, Contig1527.0.1, Contig4330.0.1, and Contig54.0.1 removed, as they are perfect duplicates of other Contigs in the assembly. Tetrahymena SMRT-seq reads were mapped to the June 2014 reference genome build. Only chromosomes with high SMRT-seq coverage (>= 80x for *Oxytricha*; >=100x for *Tetrahymena*) were used for all m^6^dA-related analyses.

### Chromosome synthesis

Synthetic Contig1781.0 chromosomes were constructed from “building blocks” of native chromosome sequence (Figures 3B, 3C, and S5). These chromosome segments were generated from either annealed synthetic oligonucleotides (with or without m^6^dA) or from genomic DNA via PCR-amplification using Phusion DNA polymerase (New England BioLabs). The latter contained terminal restriction sites for B*sa*I (New England BioLabs), a type IIS restriction enzyme that cuts distal from these sites. B*sa*I cleaves within the native DNA sequence, generating custom 4nt 5’ overhangs and releasing the non-native B*sa*I restriction site as small fragments that are subsequently purified away. The B*sa*I-generated overhangs are complementary only between adjacent building blocks, conferring specificity in ligation and minimizing undesired by-products. After B*sa*I digestion, PCR building blocks were purified by phenol:chloroform extraction and ethanol precipitation. Building blocks were then sequentially ligated to each other as described in Figure S5 using T4 DNA ligase (New England BioLabs) and purified by phenol:chloroform extraction and ethanol precipitation. Size selection after each ligation step was performed using polyethylene glycol (PEG) precipitation or Ampure XP beads (Beckman Coulter) to enrich for the large ligated product over its smaller constituents. The size of individual building blocks and their corresponding order of ligation were designed to maximize differences in size between ligated products and individual building blocks. This increases the efficiency in size selection of products over reactants.

### Verification of synthetic chromosome sequences

Synthetic chromosomes 6-8 possess 3’ single-stranded tails (Figure 3), and were first blunted by treatment with T4 DNA polymerase and DNA polymerase I, large (Klenow) fragment (New England BioLabs). All chromosomes (including 1-5) were then dA-tailed using Klenow Fragment (3’ -> 5’ exo-) (New England BioLabs), cloned using a TOPO-TA cloning kit (Thermo Fisher) or StrataClone PCR Cloning Kit (Agilent Technologies), and sequenced using flanking T7, T3, M13F, or M13R primers.

### Preparation of telomere proteins

*Oxytricha trifallax* TeBPα and TeBPβ proteins were recombinantly expressed and purified as previously described (Horvath et al., 1998).

### Preparation of *Oxytricha* histones

Vegetative *Oxytricha trifallax* strain JRB310 was cultured as previously described (Swart et al., 2013). Cells were starved for 14 hr and subsequently harvested for macronuclear isolation as previously described (Lauth et al., 1976). Purified nuclei were pelleted by centrifugation at 4000 x g, resuspended in 0.421 mL 0.4N H_2_SO_4_ per 10^6^ input cells, and nutated for 3 hr at 4°C to extract histones. Subsequently, the acid-extracted mixture was centrifuged at 21,000 x g for 15min to remove debris. Proteins were precipitated from the cleared supernatant using trichloroacetic acid (TCA), washed with cold acetone, then dried and resuspended in 2.5% (v/v) acetic acid. Individual core histone fractions were purified from crude acid-extracts using semi-preparative RP-HPLC (Vydac C18, 12 micron, 10 mM x 250 mm) with 40-65% HPLC solvent B over 50 min (Figure S4E). The identity of each purified histone fraction was verified by western blotting (Figure S4G) using antibodies: anti-H2A (Active Motif #39111), anti-H2B (Abcam #ab1790), anti-H3 (Abcam #ab1791), anti-H4 (Active Motif #39269).

### Preparation of unmodified *Xenopus* histones

All RP-HPLC analyses were performed using 0.1% TFA in water (HPLC solvent A), and 90% acetonitrile, 0.1% TFA in water (HPLC solvent B) as the mobile phases. Wild-type *Xenopus* H4, H3 C110A, H2B and H2A proteins were expressed and purified as previously described (Debelouchina et al., 2016). Purified histones were characterized by ESI-MS using a MicrOTOF-Q II ESI-Qq-TOF mass spectrometer (Bruker Daltonics). H4: calculated 11,236 Da, observed 11,236.1 Da; H3 C110A: calculated 15,239 Da, observed 15,238.7 Da; H2A: calculated 13,950 Da, observed 13,949.8 Da; H2B: calculated 13,817 Da, observed 13,816.8 Da.

### Preparation of *Xenopus* poly-acH4

H4 with N-terminal acetylation and acetylated lysines at positions 5, 8, 12, 16 and 20 was prepared by native chemical ligation. Briefly, a peptide a-thioester comprising residues 1-37 of the protein and acetylated at the appropriate positions was prepared by solid-phase peptide synthesis using the ^t^butoxycarbonyl (Boc) Nα protection strategy as described before (Fierz et al., 2011). Following cleavage and global deprotection with liq. HF, the peptide was purified by semi-preparative RP-HPLC (Vydac C18, 12 micron, 10 mM x 250 mm) using a 15-40% HPLC solvent B over 60 min, and purity was assessed by analytical RP-HPLC and ESI-MS (calculated 4278 Da, observed 4276.5 Da). H4 comprising residues 38 - 102 and an A38C mutation was prepared recombinantly with an N-terminal His-SUMO tag. Briefly, RosettaTM 2(DE3) pLysS *E. coli* cells (Novagen) were transfected with the appropriate plasmid, grown in 6 L of LB at 37 °C until 0D600 = 0.6, and induced with 1 mM IPTG for 1.5 hr. Cells were harvested by centrifugation at 5000 x g, and resuspended in lysis buffer (50 mM Tris, 150 mM NaCl, 0.1 mM EDTA, 1 mM 2-mercaptoethanol, protease inhibitors and 0.5% Triton X-100, pH 7.5). The cells were lysed by sonication, centrifuged at 30,000 x g, and the insoluble pellet was resuspended and washed in lysis buffer. The inclusion body pellet was then resuspended in 6 *M* guanidine HCl, 50 mM Tris, 150 mM NaCl, 1 mM DTT, 5 mM imidazole, pH 7.5, and the protein was bound to Ni-NTA beads at 4 ºC. The beads were washed with 6 *M* urea buffer (50 mM Tris, 1 mM DTT, 5 mM imidazole, pH 7.5), and eluted with the same buffer containing 200 mM imidazole. After purification, the protein solution was diluted to adjust the concentration of urea to 2 M, and SUMO cleavage was performed with the Ulp1 protease. The progress of the reaction was monitored by analytical RP-HPLC, and after completion, the protein solution was concentrated, incubated with 10 mM TCEP, and purified on a preparative scale by RP-HPLC and 40-60% HPLC solvent B over 60 min (calculated 7349 Da, observed 7348.3). For ligation, 2 equivalents of lyophilized a-thioester peptide (2.7 mg) were dissolved with sonication into 350 μL buffer containing 6 *M* guanidine HCl, 100 mM sodium phosphate, 200 mM 4-mercaptophenylacetic acid, pH 7.8. Meanwhile, 1 equivalent of the recombinant fragment (2.2 mg) was dissolved with sonication into 350 μL buffer containing 6 *M* guanidine HCl, 100 mM sodium phosphate, 90 mM tris(2-carboxyethyl)phosphine (TCEP), pH 7.8. The solutions were combined, and ligation was complete after 3 hr at 22.5 ºC. Purification was performed on a semi-preparative scale by RP-HPLC and a 39-57% HPLC solvent B gradient over 60 min. Pure fractions were combined and lyophilized. To convert C38 to the native alanine residue, desulfurization was performed in 6 *M* guanidine HCl, 100 mM sodium phosphate, 200 mM TCEP, 40 mM glutathione, 16 mM VA-061, pH 6.7. Briefly, 1 mg protein were dissolved in 450 μL buffer, and the reaction was complete after 16 hr incubation at 37 ºC. The final product was purified on a semi-preparative scale as described above (Figure S4). HPLC and ESI-MS equipment, and corresponding solvents were used as described for wild type *Xenopus* histone purification.

### Preparation of *Xenopus* H3K56Ac

The expression protocol was based on amber codon suppression methodology (Neumann et al., 2009), with the following modifications. The *Xenopus* H3 C110A DNA sequence (also containing the TAG codon at position 56) was cloned into a pCDF PylT1 plasmid also containing the intein MxeGyrAHis6 N198A as a C-terminal fusion. BL21(DE3) *E. coli* cells were transformed with the modified pCDF PylT1 plasmid and the pBK AcKRS3 plasmid (carrying the synthetase gene, gift from Dr. Jason Chin, MRC-LMB Cambridge), grown at 37 °C in LB media supplemented with the appropriate antibiotics (50 μg/mL kanamycin and 50 μg/mL spectinomycin). At OD600 = 0.6, 20 mM nicotinamide and 10 mM Nε-acetyl-L-lysine were added to the media. After 30 min, protein expression was induced with 1 mM IPTG for 16-18 hr. Cells were harvested by centrifugation at 5000 x g and resuspended in lysis buffer (50 mM Tris, 150 mM NaCl, 5 mM imidazole, pH 7.5), and the inclusion body pellet was isolated and resuspended in 6 *M* guanidine buffer as described above. The H3-intein fusion was purified using Ni-NTA beads (Thermo Fisher Scientific) and eluted in 6 *M* urea buffer with 500 mM imidazole. The eluted protein was dialyzed against 4 *M* urea buffer (50 mM Tris, 150 mM NaCl, 1 mM EDTA, pH 7.5) for 2 hr, and then against 2 M urea buffer for additional 2 hrs. After dialysis, 200 mM 2-mercaptoethanol were added to the solution and hydrolysis of the intein tag proceeded overnight (solution turning cloudy with precipitated H3). After hydrolysis, the concentration of urea was adjusted to 6 *M* to dissolve all protein, and 2-mercaptoethanol and EDTA were removed by dialysis against 6 *M* urea, 50 mM TrisHCl, 150 mM NaCl, pH 7.5. The hydrolyzed intein was removed by reverse Ni-NTA purification, and the H3 K56Ac protein was further purified by semi-preparative RP HPLC using a 30-70% HPLC solvent B gradient over 50 min (Figure S4).

### Preparation of histone octamers

*Oxytricha* and *Xenopus* histone octamers were respectively refolded from core histones using established protocols (Beh et al., 2015; Debelouchina et al., 2016). Briefly, lyophilized histone proteins *(Xenopus* modified or wild-type; *Oxytricha* acid-extracted) were combined in equimolar amounts in 6 M guanidine hydrochloride, 20 mM Tris pH 7.5 and the final concentration was adjusted to 1 mg/mL. The solution was dialyzed against 2 M NaCl, 10 mM Tris, 1 mM EDTA, and the octamers were purified from tetramer and dimer species using size-exclusion chromatography on a Superdex 200 10/300 column (GE Healthcare Life Sciences). The purity of the fractions was analyzed by SDS-PAGE and pure fractions were combined, concentrated and stored in 50% glycerol at -20°C.

### Quantitative mass spectrometry analysis of *Oxytricha* histone PTMs

For bottom-up MS analysis, *Oxytricha* histones were acid-extracted as described above, and dissolved in 30 μL of 50 mM NH_4_HCO_3_, pH 8.0. Derivatization reagent was prepared by mixing propionic anhydride with acetonitrile in a ratio of 1:3 (v/v), and such reagent was mixed with the histone sample in the ratio of 1:4 (v/v) for 15 minutes at 37°C. This reaction was performed twice. Histones were then digested with trypsin (enzyme:sample ratio 1:20, overnight, room temperature) in 50 mM NH_4_HCO_3_. After digestion, the derivatization reaction was performed again twice to derivatize peptide N-termini. Samples were desalted prior to nLC-MS/MS analysis by using C18 Stage-tips. Samples were analyzed by using a nLC-MS/MS setup. Chromatography was configured with the same type of column and HPLC as for the proteomics analysis. The HPLC gradient was as follows: 2% to 28% solvent B (A = 0.1% formic acid; B = 95% MeCN, 0.1% formic acid) over 45 minutes, from 28% to 80% solvent B in 5 minutes, 80% B for 10 minutes at a flow-rate of 300 nl/min. nanoLC was coupled to an LTQ-Orbitrap Elite mass spectrometer (Thermo Scientific, San Jose, CA). A full scan MS spectrum (m/z 300-1100) was acquired in the Orbitrap with a resolution of 120,000 (at 200 m/z) and an AGC target of 5x10e5. MS/MS was performed using a data-independent acquisition (DIA) mode; the entire mass range (300-1100 m/z) was fragmented at every cycle using windows of 50 m/z (16 MS/MS scans total). MS/MS AGC target was 3x10e4, the injection time limit was 50 msec and the CID collision energy was 35. MS/MS data were collected in centroid mode. EpiProfile was used to retrieve the extracted ion chromatograms and estimate the relative abundance of each peptide as compared to the total respective histone (Yuan et al., 2015). Histone protein sequence list was retrieved from the OxyDB database (http://oxy.ciliate.org/index.php/home).

Middle-down mass spectrometry analysis was performed as follows: GluC was added to the histone sample at an enzyme:sample ratio of 1:20 (overnight digestion at room temperature). Reaction was blocked by adding 1% formic acid for LC-MS analysis. 4 biological replicates were used for the analysis. Samples were separated using an Eksigent 2D+ nanoUHPLC (Eksigent, part of ABSciex). The nanoLC was equipped with a two column setup, a 2 cm pre-column (100 μm ID) packed with C18 bulk material (ReproSil, Pur C18AQ 3 μm; Dr. Maisch) and a 12 cm analytical column (75 μm ID) packed with Polycat A resin (PolyLC, Columbia, MD, 1.9 μm particles, 1000 Å). Loading buffer was 0.1% formic acid (Merck Millipore) in water. Buffer A and B were prepared as previously described (Sidoli et al., 2014). The gradient was delivered as follows: 5 min 100% buffer A, followed by a not linear gradient from 55 to 85% buffer B in 120 min and 85-100% in 10 min. Flowrate for the analysis was set to 250 nL/min. MS acquisition was performed in an Orbitrap Fusion (Thermo) with a spray voltage of 2.3 kV and a capillary temperature of 275 °C. Data acquisition was performed in the Orbitrap for both precursor and product ions, with a mass resolution of 60,000 for MS and 30,000 for MS/MS. MS acquisition window was set at 660-740 m/z. Dynamic exclusion was disabled. Precursor charges accepted for MS/MS fragmentation were 5-8. Isolation width was set at 2 m/z. The 5 most intense ions with MS signal higher than 5,000 counts were isolated for fragmentation using ETD with an activation time of 20 msec. 3 microscans were used for each MS/MS spectrum, and the AGC target was set to 2x10e5. Data processing was performed as previously described (Sidoli et al., 2014). Briefly, spectra were deconvoluted with Xtract (Thermo) and searched with Mascot (v2.5, Matrix Science, London, UK), including acetylation (K) as dynamic modifications. No fixed modifications were selected. Histone protein sequence list was retrieved from the OxyDB database (http://oxy.ciliate.org/index.php/home). Enzyme was GluC (cleaves after E) with 0 missed cleavages allowed. Mass tolerance was set to 2.1 Da for precursor mass and 0.01 Da for product mass. Mascot result files were processed using a tolerance of 30 ppm, as we previously demonstrated it is a suitable value to filter confident identification and quantification (Sidoli et al., 2014). Peptides with ambiguous modification site assignments were automatically discarded by the software.

Middle-down and bottom-up MS-analysis were performed with four biological replicates (n_b_ = 4). Moreover, each biological replicate of the histone sample was performed in technical triplicates (n_t_ = 3) (with the exception of “Replicate 4” for middle-down data, where n_t_ = 2). No samples were excluded as outliers (this applies to all proteomics analyses described in this manuscript).

### Preparation of mini-genome DNA

98 full-length chromosomes were individually amplified from *Oxytricha trifallax* strain JRB310 genomic DNA using Phusion DNA polymerase (New England BioLabs). Primer pairs are listed in Table S2. Amplified chromosomes were separately purified using a MinElute PCR purification kit (QIAGEN), and then mixed in equimolar ratios to obtain “mini-genome” DNA. The sample was concentrated by ethanol precipitation and adjusted to a final concentration of ~1.6mg/ml.

### Preparation of native genomic DNA for chromatin assembly

Macronuclei were isolated from vegetative *Oxytricha trifallax* strain JRB310 as previously described (Lauth et al., 1976), and genomic DNA was purified using the Nucleospin Tissue kit (Macherey-Nagel). Approximately 200 μg of genomic DNA was loaded on a 15%-40% linear sucrose gradient and centrifuged in a SW40Ti rotor (Beckman Coulter) at 160,070 x g for 22.5hr at 20°C. Sucrose solutions were in 1 *M* NaCl, 20 mM Tris pH 7.5, 5 mM EDTA. Individual fractions from the sucrose gradient were analyzed on 0.9% agarose-TAE gels. Fractions containing high molecular weight DNA that migrated at the mobility limit were discarded as such DNA species were found to interfere with downstream chromatin assembly. All other fractions were pooled, ethanol precipitated, and adjusted to 0.5mg/mL DNA.

### Telomere protein binding to synthetic chromosomes

Synthetic chromosomes were mixed with a 100-fold excess of non-specific competitor DNA (full length Contig20883.0 DNA lacking telomeric repeats), prepared by PCR amplification from genomic DNA and purified with Ampure XP beads (Beckman Coulter). 1 μg competitor DNA was mixed with 0.9 *μM* TeBPα and 0.9 *μM* TeBPβ proteins in a 10 μL volume containing 10 mM Tris pH 7.5, 0.1 mM EDTA at room temperature for 1 hr. Subsequently, 15 ng synthetic chromosome was added and incubated at room temperature for 1 hr. To assess TeBP binding to chromosome ends, an aliquot of DNA : TeBPα : TeBPβ complex was adjusted to 50 mM NaCl, 100 mM Tris pH 7.5, 10 mM MgCl_2_, 0.025% (v/v) Triton X-100 and digested with 0.625 Units *SspI* (New England BioLabs) for 1 hr at 37°C before transferring to ice. An aliquot of each reaction was adjusted to 2 M NaCl to challenge protein binding with high salt. All samples were loaded on native 5% polyacrylamide gels in 0.5x TBE at 4°C to assess TeBP binding to chromosome ends. The remaining undigested DNA : TeBPα : TeBPβ complexes were used for chromatin assembly by salt gradient dialysis as described below.

### Chromatin assembly and preparation of mononucleosomal DNA

All chromatin assemblies were prepared by salt gradient dialysis as previously described (Beh et al., 2015; Luger et al., 1999). To reduce sample requirements while maintaining adequate DNA concentrations for chromatin assembly, synthetic chromosomes were first mixed with a hundred-fold excess of “carrier” DNA (PCR-amplified *Oxytricha* Contig17535.0). We verified that nucleosome occupancy in terminal regions (qPCR primer pairs 1 and 24) and the methylated region (qPCR primer pairs 6 and 7) of the synthetic chromosome is unaffected by the presence of carrier DNA (Figure S8). Native and mini-genome DNA were not mixed with carrier DNA prior to chromatin assembly.

Histone octamers and (synthetic chromosome + carrier) DNA were mixed in a 0.8:1 mass ratio, while histone octamers and (native or mini-genome) DNA were mixed in a 1.3:1 mass ratio, each in a 50 μL total volume. Samples were then dialyzed into start buffer (10 mM Tris pH 7.5, 1.4 *M* KCl, 0.1 mM EDTA pH 7.5, 1 mM DTT) for 1 hr at 4°C. Then, 350 mL end buffer (10 mM Tris pH 7.5, 10 mM KCl, 0.1 mM EDTA, 1 mM DTT) was added at a rate of 1 mL/min with stirring. The assembled chromatin was then dialyzed overnight at 4°C into 200 mL end buffer, followed by a final round of dialysis in fresh 200 mL end buffer for 1 hr at 4°C. The assembled chromatin was then adjusted to 50 mM Tris pH 7.9, 5 mM CaCl_2_ and digested with MNase (New England BioLabs) to mainly mononucleosomal DNA as previously described (Beh et al., 2015). Mononucleosome-sized DNA was gel-purified and used for tiling qPCR on a Viia 7 Real-Time PCR System, or *in vitro* MNase-seq on an Illumina HiSeq 2500, according to the manufacturer’s instructions. qPCR primer sequences are listed in Table S2.

### Tiling qPCR analysis

qPCR data were analyzed using the ΔΔCt method (Livak and Schmittgen, 2001). For each qPCR locus, the Ct value generated from mononucleosomal DNA was normalized to data from the corresponding naked, undigested synthetic chromosome. This controls for potential variation in PCR amplification efficiency, especially over methylated regions. The calculated fold change in qPCR signal in nucleosomal DNA, relative to the naked chromosomal DNA control, is denoted as ‘nucleosome occupancy’ for all presented qPCR data. Primer pair 22 was used as the reference for methylated chromosome experiments, while primer pair 12 was used for TeBP binding experiments (see Figure 4 and Table S2 for primer positions and sequences). These primer pairs are distant from the sites of DNA methylation and TeBP binding, respectively.

### ACF spacing assay

ATP-dependent nucleosome spacing was performed in accordance with a previous study(Lieleg et al., 2015). Chromatin was assembled by salt gradient dialysis as described above, and then adjusted to 20 mM HEPES-KOH pH 7.5, 80 mM KCl, 0.5 mM EGTA, 12% (v/v) glycerol, 10 mM (NH_4_)_2_SO_4_, 2.5 mM DTT. Samples were then incubated for 2.5 hr at 27°C with 3 mM ATP, 30 mM creatine phosphate, 4 mM MgCl_2_, 5 ng/μL creatine kinase, and 11 ng/μL *Drosophila* ACF (purchased from Active Motif). Remodeled chromatin was then adjusted to 5 mM CaCl_2_, and subject to MNase digestion, mononucleosomal DNA purification, and qPCR analysis as described above.

### Quantification and statistical analysis

All statistical tests were performed in R (v3.2.5), and described in the respective Figure and Table legends.

### Data availability

All raw files from histone mass spectrometry are deposited in the CHORUS database under project number 1298 (https://chorusproject.org/). *Oxytricha* SMRT-seq data are deposited in SRA under the accession numbers SRX2335608 and SRX2335607. *Tetrahymena* SMRT-seq and all *Oxytricha* Illumina data are deposited in NCBI GEO under accession number GSE94421.

